# Oscillatory Entrainment Guides Temporal Prediction in Non-Deterministic Continuous Environments, Independent of Bayesian Sequential Learning

**DOI:** 10.64898/2026.11.11.765929

**Authors:** Elmira Hosseini, Assaf Breska

**Affiliations:** Max Planck Institute for Biological Cybernetics, Tübingen, Germany; Graduate Training Centre of Neuroscience, University of Tübingen, Tübingen, Germany

## Abstract

Predicting future event timing is essential for human interaction with our environment, and has been established in stimulus streams containing deterministic regularities. However, real-world streams often lack such regularities, leaving it unclear whether and how temporal predictions operate. Here we show that oscillatory entrainment, a mechanism identified for deterministic streams, operates in non-deterministic streams containing substantial temporal jitter. Model simulations revealed divergent dynamics and predicted timepoints between the oscillatory mechanism, which tracked local input regularities, and Bayesian learning, an alternative computational mechanism, which encoded global statistics. Across two speeded detection experiments using mechanism-dissociating streams, human performance was facilitated for targets appearing at oscillator-predicted times in streams with intermediate jitter levels, relative to maximal-jitter streams. Importantly, this effect was decoupled from Bayesian-based prediction and non-specific preparation, and could flexibly adapt to stream rate variations. These findings reveal the parallel mechanisms employed by humans when faced with continuous temporal uncertainty.

## Main

Adaptive behavior critically relies on the ability to anticipate not only what will occur where but also when, allowing organisms to proactively optimize perception and action^1–4^. Research has shown that humans can form temporal predictions based on deterministic temporal regularities. When a known interval separates two isolated events, such as between the traffic light changing from red-yellow to green, predictions rely on dedicated interval timing circuits that compare elapsed time from the first event to the memorized interval^5,6^. Conversely, when a continuous stream is isochronous, such as beat in music, predictions rely on entrainment (coupling) of endogenous neural oscillations, aligning the optimal phase for performance to the beat^2,7^. However, while natural sensory streams are continuous, they rarely adhere to strictly deterministic temporal structures (e.g., speech or natural motion). Yet, how temporal predictions operate in environments comprised of continuous stream of events with variable intervals in between, remains unknown.

Previous research on temporal prediction under uncertainty mostly focused on cases where isolated cue-target intervals vary across trials. These studies found that behavior and neural dynamics reflected the statistical properties of interval distributions^8–10^, later attributed to Bayesian sequential updating mechanisms^10–12^. Hypothetically, a similar mechanism could operate in continuous non-deterministic environments, with intervals integrated sequentially. Moreover, oscillatory mechanisms could still operate due to the continuous environment, leading to benefits at optimal phases^13^. To date, it is unclear whether humans use either of these mechanisms when interacting with continuous jittered environments, or whether no specific predictions are formed (‘vigilant mode’)^14^, or anticipation is increased with elapsed time^8^.

Addressing this fundamental gap faces multiple challenges. Computationally, it is unknown under what conditions a fixed-period coupled oscillator mechanism, previously shown to explain prediction in isochronous streams^15^, can systematically guide predictions in jittered streams. Whereas in isochronous streams the optimal oscillatory phase aligns to on-beat time independently of the initial state (angle) of the oscillator, in jittered streams the specific structure could interact with the initial state to lead to phase dispersion, precluding its usability to guide prediction^16^. Second, even if an oscillator-based mechanism could lead to consistent prediction, it is not clear how this is dissociated from Bayesian mechanisms.

Recent studies suggested that an oscillator mechanism could depend on Bayesian processes or approximate Bayesian updating of phase or rate in some jittered environments^13,17^, but it is unknown how they dissociate in less constrained streams akin to our rich natural environment. Moreover, such mechanism is less parsimonious than a fixed-period oscillator^18^, and to date, the segregation between oscillator and Bayesian interval learning processes remains highly debated^19^. Finally, it is unclear whether humans leverage these computational principles in high-jitter environments to drive prediction. Studying high-jitter environments is of special interest as they better resemble naturalistic environments, such as speech^20,21^.

Here, we combined computational modelling and model-based behavioral experiments to disentangle oscillatory entrainment and sequential Bayesian learning accounts of temporal prediction in continuous sensory streams with temporal variability. Using large-scale model simulations, we asked whether a fixed-period coupled oscillator mechanism could lead to specific predictions in temporally variable streams, whether and when its predictions and representation of temporal uncertainty can be dissociated from those of a Bayesian learning process, and how this dissociation depends on both environmental statistics and internal model parameters. We then examined how these computational strategies accounted for human behavior, in two experiments that required speeded detection of targets embedded in continuous temporally jittered streams with varying degrees of interval variability. Critically, the modelling results were used to guide the streams’ design to decouple the predicted time points of the two processes, allowing us to isolate predictive benefits obtained by each of the two mechanisms separately.

## Results

### Computational Modelling and Simulations

To investigate how Oscillator and Bayesian mechanisms could support temporal prediction in non-deterministic continuous stream of events, we first implemented a computational modelling framework to identify overlapping or distinguishable model behaviors. The oscillatory entrainment process was modelled using a fixed-period coupled oscillator (hereafter Oscillator model), a model previously shown to parsimoniously explain temporal predictions in isochronous streams^22^. The Bayesian sequential updating process was modelled using a conjugate prior ideal Bayesian observer (hereafter Bayesian model), an extended version of a model used previously to explain temporal predictions formed across variable isolated intervals^10^. We tested these two models on a large cohort of stimulus streams with varying degree of inter-stimulus interval (ISI) variability, and evaluated their behavior through examining the time points they predicted and the change in their prediction uncertainty relative to an uninformed state.

### Oscillator and Bayesian model predictions can be separated despite general overlap

We first asked whether a fixed-period oscillator can generate stable temporal predictions in a jittered environment and, if so, whether these predictions differ from those of a Bayesian model (Fig. 1a). In streams in which the ISI jitter is 25% of the mean (medium variance relative to a corresponding uniform distribution), the Bayesian model generated predicted points that converged to the stream mean across different streams, whereas the Oscillator model generated a broader spread of predicted points with varying levels of uncertainty reduction (measured as phase concentration, PC). Despite temporal variability, many streams produced robust PC (32.3% above threshold), attesting to stable Oscillator predictions. Importantly, some of these streams (3.3%) also exhibited near-maximal separation between the Oscillator and Bayesian predictions (narrow window around ±*π*; shaded regions, Fig. 1a, for example stream see Fig. 1b). Thus, reliable Oscillator-based predictions can be separated from Bayesian-based predictions.

**Figure 1.**
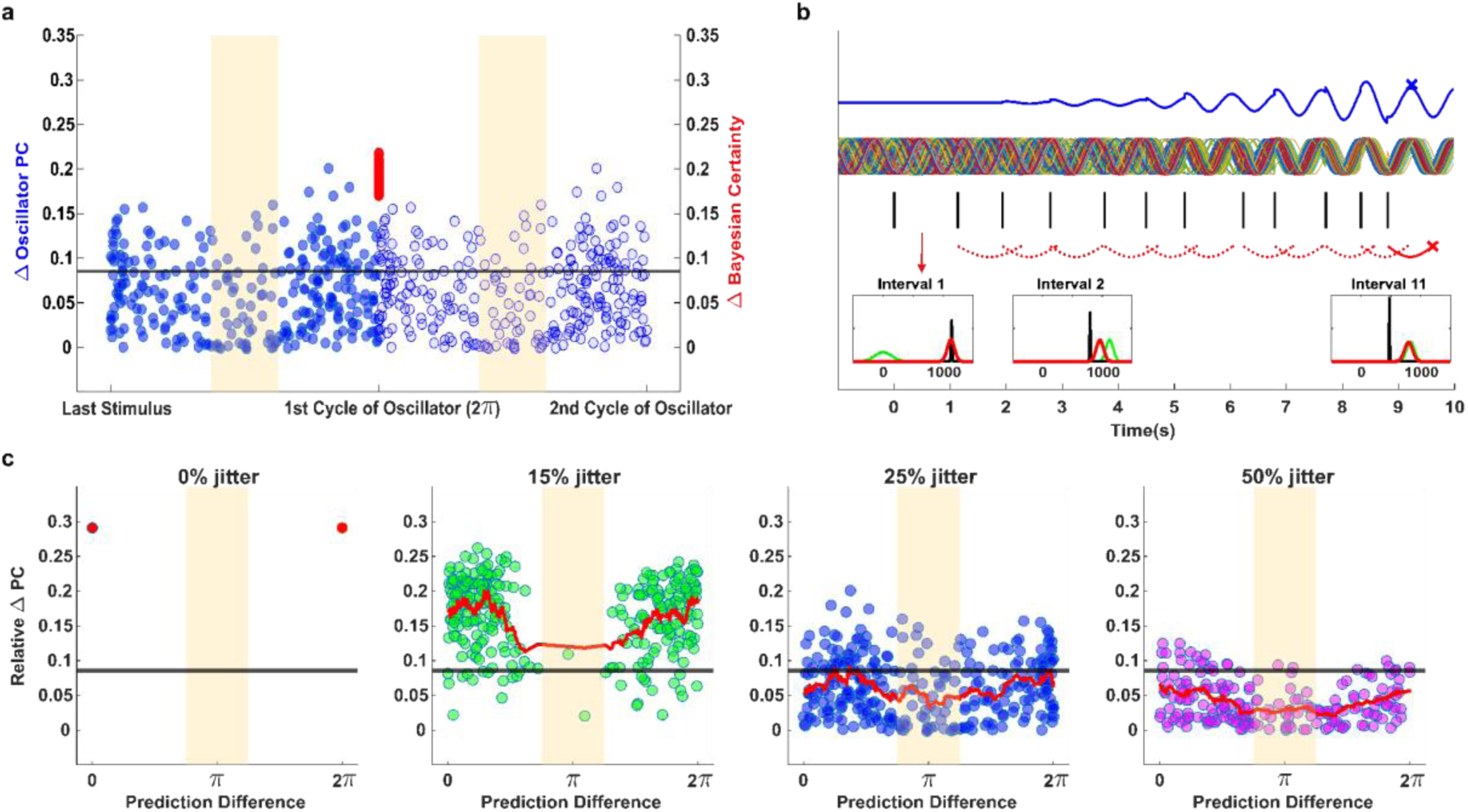
Predictions of the Oscillator and Bayesian models across temporal variability. **a.** Predicted time points of the models and their certainty levels. Filled dots represent different shuffles of the ISIs of a stimulus stream with 25% jitter. Blue: Oscillator model; red: Bayesian model; Both colors are the same set of shuffles. For this visualization, relative Bayesian certainty was obtained by 1-*abs*(Δ*uncertainty*). Transparent blue dots represent the second cycle prediction point of the Oscillator model in the same set of shuffles. Shaded region: predefined window centered around half cycle of the Oscillator model (here equivalent to π), where model predictions are most differentiated. Black horizontal line: PC significance threshold. **b.** Oscillator and Bayesian models’ dynamics in response to a 25% jittered stream, which leads to high prediction difference. Top: Oscillator model single trial traces (multicolor) and predicted angle weighted by certainty (blue), with final predicted time (x) at optimal phase. Middle: stimulus onset times (black). Bottom: Predicted time points of Bayesian model after each stream interval (red), with final predicted time (x), and illustrations of prior (green), likelihood (black) and posterior (red) for three steps. The Oscillator model converges to a stable prediction, the Bayesian model converges to predicting the mean of the stream ISIs. **c.** Prediction difference between the two models and prediction certainty of Oscillator for different stream jitter levels. Left to right: 0% (isochronous), 15%, 25%, and 50% ISI variability. Each dot represents a specific shuffle of the ISIs of a stream with defined mean and variance. The red line is the moving average of relative PC change across shuffles. The 0% (isochronous) condition only shows two single dots at the first and second cycle of the Oscillator as shuffles are identical.

We next examined how this separation changes as temporal variability increases (Fig. 1c). In isochronous streams, the predictions of both models completely overlapped, converging on the on-beat time corresponding to the mean stream interval. As temporal variability increased (0%, 15%, 25%, 50% jitter), there were more streams showing divergence between the predictions of the two models, despite all being different shuffles of the same ISI set at each variability level, hence having identical global stream statistics (0%, 0.6%, 17.6%, 16% of shuffles in the shaded region, respectively). However, larger ISI variabilities were accompanied by progressively lower PC (100%, 90.6%, 32%, 13% above chance threshold, respectively). This trade-off between the separability of Oscillator and Bayesian predictions and the stability of oscillatory entrainment led to a non-linear relationship between ISI variability and the presence of shuffles with above threshold PC and in shaded region (0%, 0.3%, 3.3%, 1%, respectively). Thus, crucially, although Oscillator and Bayesian predictions generally overlap across a large set of jittered environments^23^, they can nevertheless generate distinct and reliable predictions in specific streams of events with medium ISI variability.

### Temporal variability induces divergence in uncertainty representation between Oscillator and Bayesian models

To investigate the source of the divergence between the two models, we compared their representation of temporal uncertainty in two analyses. In the first analysis, we directly compared the final model uncertainties (in Oscillator model defined as phase variance, PV) across a broad range of stream ISI variability, within each of which across a cohort of shuffles of the same set of ISIs (Fig. 2a). An overlap hypothesis would predict a linear relationship between the uncertainty of the two models across levels of stream variabilities. However, the relationship between the two model uncertainties was nonlinear (ΔAIC=27.6 relative to a linear model), with uncertainty in the Bayesian model increasing monotonically with temporal variability, but in the Oscillator model stabilizing after 40% jitter (Fig. 2a). To verify that this nonlinear relationship was not an outcome of the uncertainty measures used (e.g., circular vs. linear), we analyzed the models in isochronous streams, where model overlap is expected, manipulating input certainty by increasing stream length (Fig. 2b). Indeed, the degree of model uncertainty was linearly related between the Bayesian and Oscillator models (ΔAIC=3.7 in favor of linear model). Thus, the nonlinear relationship observed with increasing temporal variability is not a consequence of the different measures, but rather reflects true divergence in how the models represent uncertainty.

**Figure 2.**
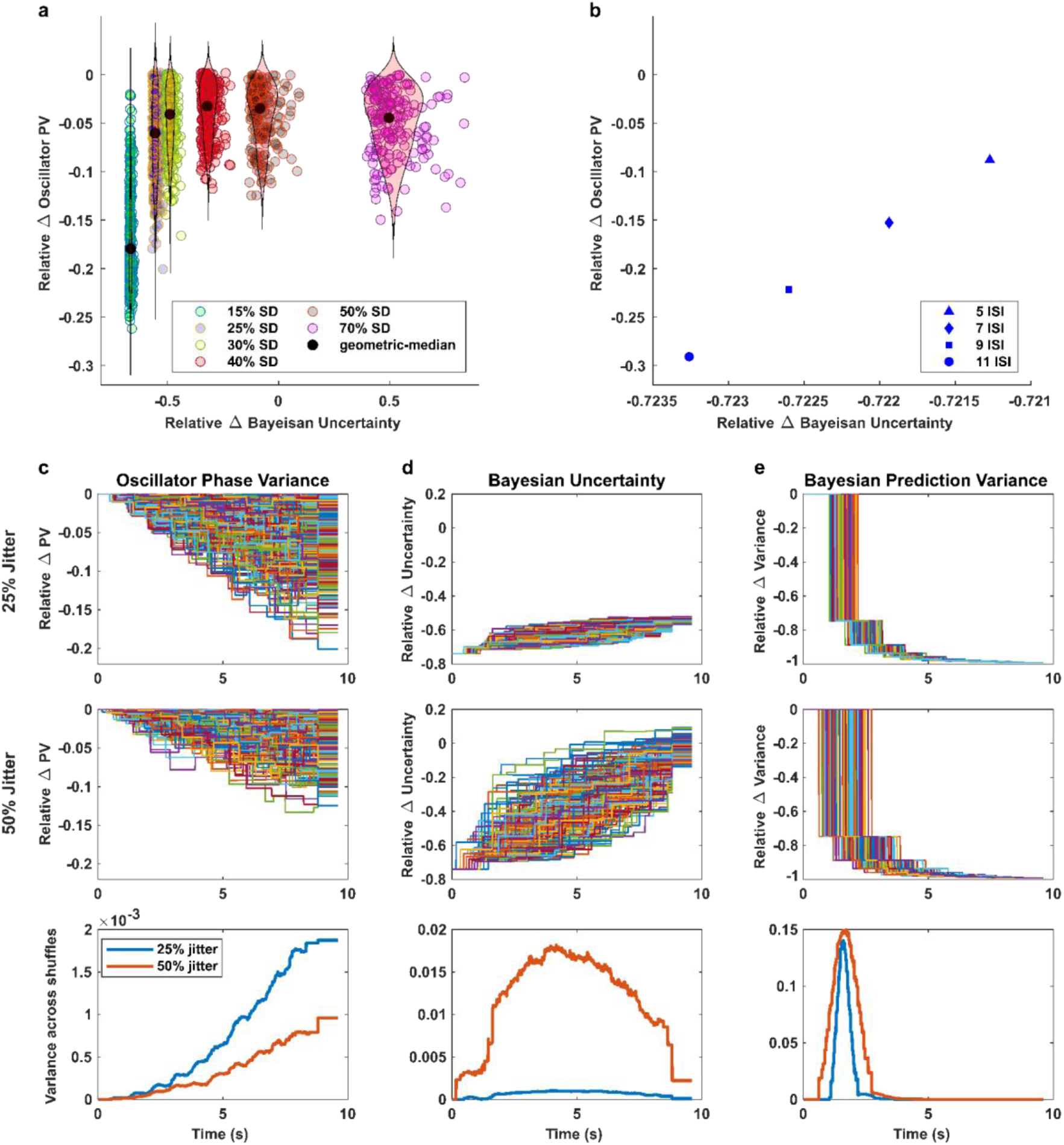
Uncertainty of the Oscillator and Bayesian models across jittered and isochronous input streams. **a.** Relative uncertainty change compared to a uniform distribution by the Bayesian and the Oscillator models after the final stream stimulus, for streams with ISI variability ranging from 15–70% (SD). Each dot represents one shuffle; black markers indicate the geometric median. **b.** Same as a. for perfectly isochronous streams, for different input lengths. **c.** Trajectory of Oscillator model uncertainty throughout the stream. Top: time-resolved dynamics of relative PV decrease (quantified as 1-PC) compared to the same measure in a uniform distribution, across different shuffles of streams with 25% jitter. Middle: same for streams with 50% jitter, Bottom: variance in trajectory dynamics across shuffles of each jitter condition. **d.** Trajectory of Bayesian model uncertainty throughout the stream. Same as c., with decrease of the uncertainty (quantified as mean of the inverse-gamma distribution) relative to a uniform distribution. Note that in top and middle panels, the starting point, and hence increasing trend, is based on an arbitrarily chosen uncertainty around each uniformly distributed prior values. **e.** Trajectory of Bayesian model Prediction Variance throughout the stream. Same as d., but with decrease of the variance of the predictions generated from uniformly distributed priors standardized to the variance after first interval.

In the second analysis, we examined the only trajectory of prediction uncertainties (as it is not possible to reliably track the trajectory of predicted time points themselves for the Oscillator model, see Methods) in the Oscillator and Bayesian models as the stream unfolds, and how it is modulated by ISI variability (Fig. 2c-e). We asked whether the dynamics of model uncertainties overlap. For the Oscillator model (Fig. 2c), there were markedly divergent trajectories of uncertainty reduction and final levels of PV for 25% jittered streams, even though they were different shuffles of the same ISI set, with this divergence increasing as the stream progressed (Fig. 2c, top). This pattern was present but weaker for 50% jittered streams, due to weaker stabilization of predictions (Fig. 2c, middle) and therefore less uncertainty reductions. This led to higher PV variability across all the streams of 25% compared to 50% jitter (Fig. 2c, bottom).

In contrast, the Bayesian model, exhibited a qualitatively different pattern. The trajectories across streams initially diverged, but then converged to a specific level, that was expectedly lower in the 25% jitter condition (Fig. 2d, top) than the 50% condition (Fig. 2d, middle). Moreover, the initial divergence was markedly larger in 50% jittered condition, contrary to the Oscillator model (Fig. 2d, bottom). To ensure this discrepancy is not measure-specific, we also evaluated the Bayesian model by computing an alternative uncertainty metric, Prediction Variance (see Methods, Fig. 2e). Here near-complete reduction occurred both in 25% and 50% jitter conditions (Fig. 2e, top, middle), indicating robust convergence of predictions themselves even under elevated temporal variability. Moreover, this was highly consistent across different streams and jitter levels (Fig. 2e, bottom). These results indicate that the Oscillator model is sensitive to the specific temporal structure of the input stream, whereas the Bayesian model consistently converges to estimate the statistics of the distribution, regardless of temporal ordering of intervals.

### Variations in parameter-controlled input sensitivity and forgetting reveal differential balancing of local and global regularities

As a complementary analysis, to understand the representation of uncertainty in Oscillator and Bayesian models, we examined their behavior while systematically varying their input sensitivity and forgetting, via model-specific parameters. Rather than a technical aspect of parameter selection, we aimed to gain insight into the relationship between these two mechanisms based on their susceptibility to functionally analogous parameters. Importantly, we examined the transition from isochronous to temporally jittered environments, hypothesizing that this could reveal distinct patterns. For isochronous streams we quantified relative uncertainty reduction across different stream lengths, whereas for jittered streams, across different shuffles of the ISIs of a 25% jittered stream.

We first examined variations in the input sensitivity parameter (Oscillator: entrainment strength, Bayesian: prior dependency). For the Oscillator model (Fig. 3a, left), in isochronous streams, stronger input sensitivity produced monotonic reduction in uncertainty (lower PV) within each stream length (Fig. 3a, top left), making stronger input sensitivity seem invariantly beneficial. In jittered streams however, while this pattern was preserved on average, it strongly depended on the precise temporal arrangement of intervals, with some shuffles showing no reduction in uncertainty even with strong entrainment (Fig. 3a, bottom left). For the Bayesian model (Fig. 3a, right), in isochronous streams, higher input sensitivity (lower prior dependency) led to monotonically lower uncertainty, similar to the Oscillator model. Strikingly, in jittered streams (Fig. 3a, bottom right) this pattern was preserved, with much smaller variability between shuffles within each input sensitivity level compared to the Oscillator model. Thus, the Bayesian model did not show sensitivity to the specific interval order, as shown by the Oscillator model.

**Figure 3.**
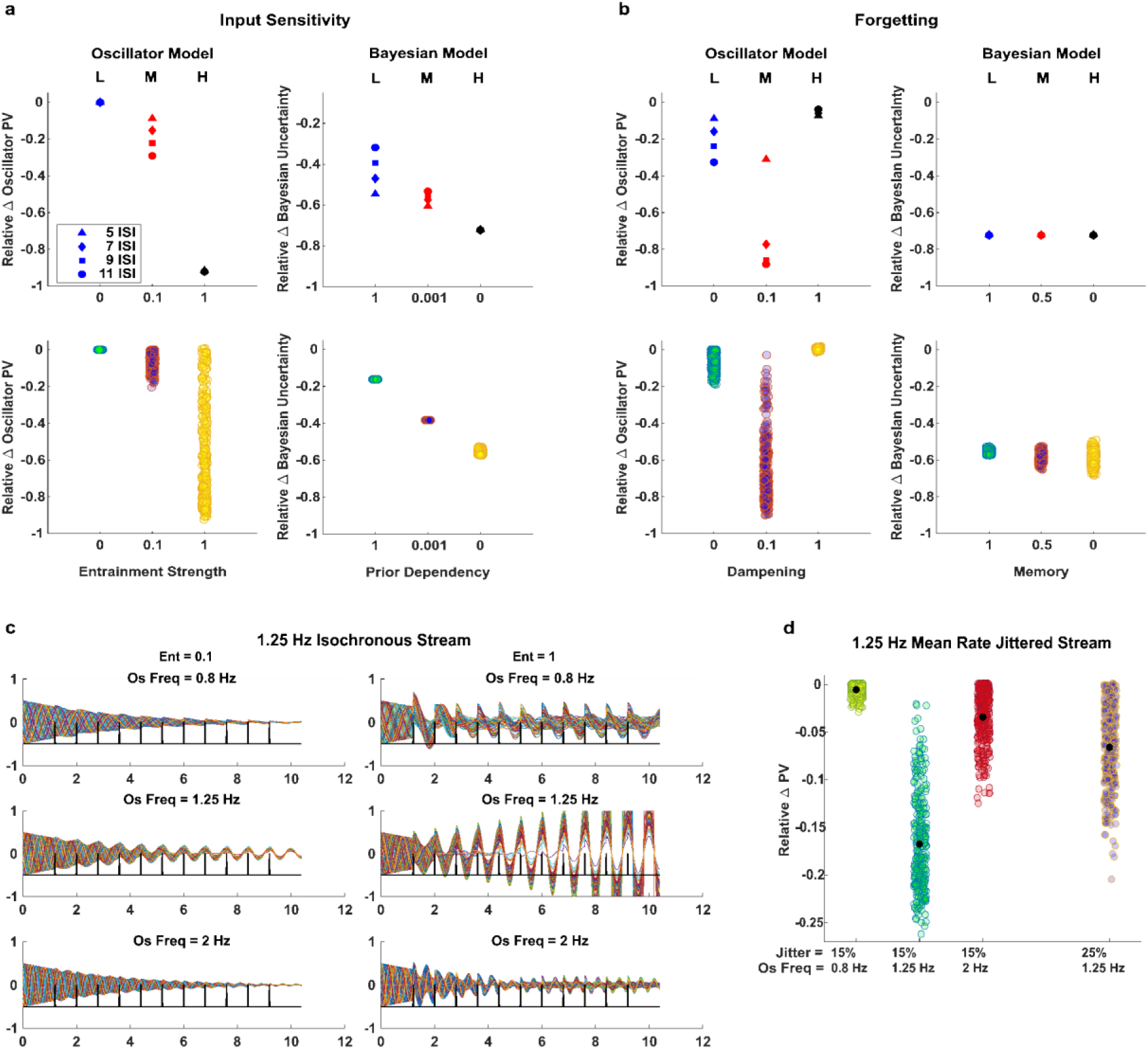
Effects of variation in internal model parameters on uncertainty reduction across isochronous and jittered input streams. **a.** Impact of varying Input sensitivity of Oscillator (left) and Bayesian (right) models across isochronous (top row; 5–11 intervals) and jittered streams (bottom row; 25% ISI variability, 11 intervals; Each dot represents a shuffle of the input stream) on uncertainty reduction. Input sensitivity was set to High, Medium or Low (H, M, L, respectively), and manipulated in Oscillator via entrainment strength (c = 0, 0.1, 1) and in Bayesian via prior dependency (κ₀ = 0, 0.001, 1). **b.** Same as a., for varying Forgetting, manipulated in Oscillator via dampening strength (d = 0, 0.5, 1), and in Bayesian by introducing a memory factor (mf = 0, 0.5, 1). **c.** Time courses of dampening oscillators starting from randomly distributed initial angles for 1.25 Hz isochronous input with input-matching and-mismatching oscillator frequencies and different entrainment strengths. **d.** Relative uncertainty reduction (PV change) in Oscillator model for a jittered stream with 1.25 Hz mean-rate, with varying jitter levels and oscillator frequencies. Black dots represent condition means; colored dots represent individual shuffles.

Variation in the forgetting parameter (Oscillator: dampening, Bayesian: memory, Fig. 3b) showed a similar pattern. For the Oscillator model, optimal dampening level (medium) led to strong uncertainty reduction in isochronous streams. However, in jittered streams, while this pattern was observed on average it highly depended on the temporal configuration of individual streams, with some shuffles showing no uncertainty reduction even in medium forgetting (Fig. 3b, bottom left). For the Bayesian model, strong uncertainty reduction was observed in all forgetting levels in isochronous streams (Fig. 3b, top right; note no effect of forgetting in isochronous streams, likely due to setting input sensitivity to high, see Methods). Most importantly, strong reduction was preserved across forgetting levels in all shuffles of jittered streams, with smaller spread across shuffles compared to the Oscillator model (Fig. 3b, bottom right). Together, these results highlight a fundamental computational disparity between the two models: while in the Oscillator model uncertainty reflected sensitivity to local temporal structure, Bayesian uncertainty was predominantly governed by the global statistics and how evidence was statistically weighted and accumulated over time.

Finally, as we did not use an adaptive frequency oscillator model, we examined the impact of frequency match between oscillator and input stream, asking whether the superiority of a frequency-matching oscillator generalizes to the jittered streams. For isochronous streams and intermediate input sensitivity (entrainment strength), a frequency-matched oscillator exhibits stronger PC and sustained amplitude relative to oscillators tuned ±0.5 octave away from the input (Fig. 3c, Left; cf.^16,22^). While, as the results above (Fig. 3a) suggest, high entrainment strength is beneficial in driving stronger reduction of uncertainty, this comes at a cost of global selectivity, as multiple oscillators exhibited high PC and sustained amplitude (Arnold tongue-like pattern, Fig. 3c, Right). We then examined the impact of temporal jitter (15%), which could hypothetically eliminate frequency specificity due to specific intervals aligning mismatched oscillators. Surprisingly, the frequency-matched oscillator still exhibited the strongest PV reduction (Fig. 3d), and mismatched oscillators performed worse even than the frequency-matched oscillator run on a more jittered stream (25%). Thus, under intermediate entrainment strength a frequency matching oscillator stands out even under higher jitter, in its ability to robustly guide prediction.

### Behavioral Experiments

Given our simulations’ findings that Oscillator and Bayesian predictions can be dissociated in jittered streams, we turned to examine how humans used these two mechanisms in two behavioral experiments. We selected a subset of streams with 25% and 50% ISI variability and various lengths that produced the maximal dissociation of models’ predictions while, in the 25% condition, still yielding sufficient phase-locking in the Oscillator model to produce interpretable predictions. These were used in speeded detection experiments (Fig. 4a) in which targets could coincide, with equal probability, with the Oscillator prediction, Bayesian prediction, or two additional time points as control (Fig. 4b).

**Figure 4.**
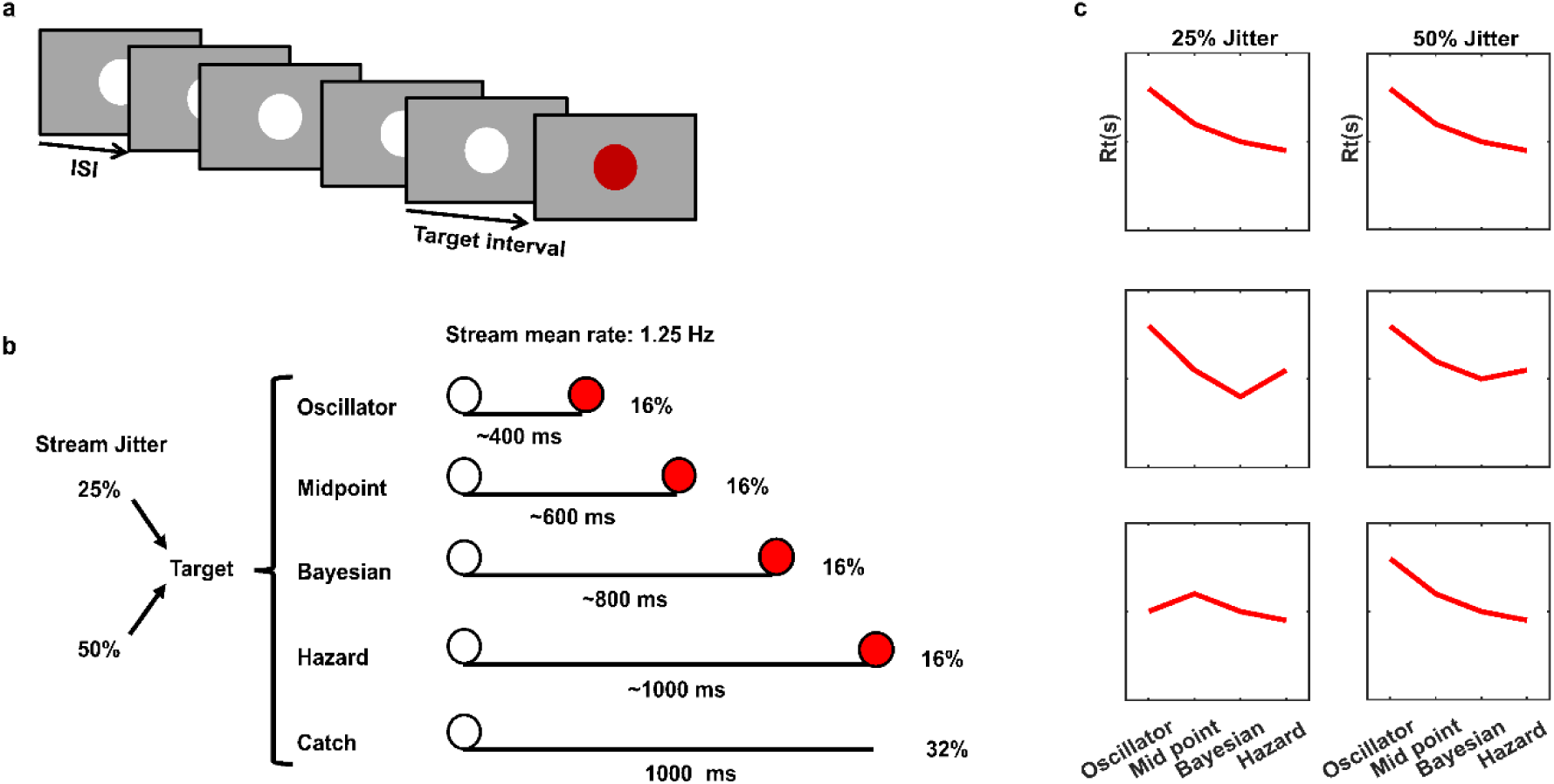
Behavioral paradigm and different hypotheses for response time patterns. **a.** Trial structure. A sequence of white visual flashes, with the specific ISI order determined based on modelling results to dissociate model predicted times, followed by a red target stimulus after a controlled target interval. **b.** Experimental design: target intervals match the predictions of the Oscillator or Bayesian models, or two additional control time points (midpoint, hazard rate). Percentages indicate the relative amount of each target interval condition or of target-absent trials (Catch) within a block. The illustrated target intervals are for a stream with mean rate of 1.25 Hz (800 ms mean interval) used in Experiment 1 **c.** Three non-mutually-exclusive hypothesized patterns of mean reaction times (RTs) to the four target interval conditions, separately for 25% jitter (left column) and 50% jitter (right column). Top: Hazard-only hypothesis. RTs decrease monotonically with later targets in both levels of jitter due to increasing conditional probability of target occurrence. Middle: Bayesian hypothesis. In addition to a hazard effect, RTs are selectively facilitated for targets appearing at Bayesian-predicted interval, reflecting sequential updating as a prediction mechanism. If the strength of Bayesian prediction reflects certainty, this should be stronger in medium jitter condition (25%). Bottom: Oscillator hypothesis. In addition to a hazard effect, RTs are selectively facilitated for targets appearing at Oscillator-predicted interval for the comparably lower jitter (25%), reflecting entrainment-based temporal prediction.

### Behavioral facilitation for Oscillator predicted events, decoupled from Bayesian predictions

The first experiment (N=26) included streams with a mean rate of 1.25Hz (mean interval 800ms), with three ISI variability conditions, SD=25% (medium variance, MV), 50% (high variance, HV) and 0 (isochronous) in separate blocks. We considered three non-mutually exclusive hypotheses (Fig. 4c): Hazard-effect: increased preparation with the passage of time should decrease reaction times (RTs) for later targets; Bayesian-based prediction: faster RTs to the Bayesian targets relative to control targets, and possibly in MV relative to HV due to the lower uncertainty; Oscillator-based prediction: faster RTs to the Oscillator targets in MV as these had robust PC, relative to the middle targets and/or to the HV due to the decrease in uncertainty.

For the jittered conditions (Fig. 5a), RTs were overall faster in the MV compared to HV condition (2×4 repeated-measures ANOVA, main effect of jitter: F(1,25)=23.48, p<.001, η²p=.48). Also, RTs varied between target positions (F(3,75)=10.16, p<.001, η²p=.29), becoming progressively faster for later targets, consistent with a hazard-rate effect. Critically, the interaction between jitter and target position was significant (F(3,75)=6.48, p<.001, η²p=.21), indicating that temporal variability differentially affected performance across prediction time points. Directly assessing the impact of temporal variability on model-predicted target times revealed faster RTs in the MV condition for Oscillator-predicted targets (t(25)=−7.36, p<.001, d=1.44) and for Bayesian-predicted targets, although with a smaller effect (t(25)=−2.85, p=.009, d=.56). To evaluate these effects beyond overall condition differences, we repeated these comparisons after subtracting the hazard control target RT in each condition. This revealed faster RTs in the MV condition for Oscillator-predicted targets (t(25)=−2.48, p=.02, d=.49), but not for Bayesian-predicted targets (trend in the opposite direction, t(25)=1.60, p=.12, d=.31). Thus, Oscillator-based predictions in streams with medium temporal variability benefit performance beyond the overall speeding relative to high temporal variability, whereas Bayesian predictions largely tracked general changes in response speed. Finally, directly comparing the impact of reduced jitter on Oscillator-and Bayesian-based targets revealed that the RT reduction for Oscillator-predicted targets was greater than that for Bayesian-predicted targets (2×2 repeated-measures ANOVA, Jitter×Model interaction: F(1,25)=14.68, p<.001, η²p=.37). There was also a significant main effect of prediction model, F(1,25)=17.20, p<.001, η²p=.41, but no main effect of jitter, F(1,25)=.70, p=.41, η²p=.03. Together, these results demonstrate that Oscillator-based predictions operate in aperiodic streams, producing a stronger behavioral benefit under more predictable temporal conditions, even when decoupled from the prediction of a Bayesian mechanism.

**Figure 5.**
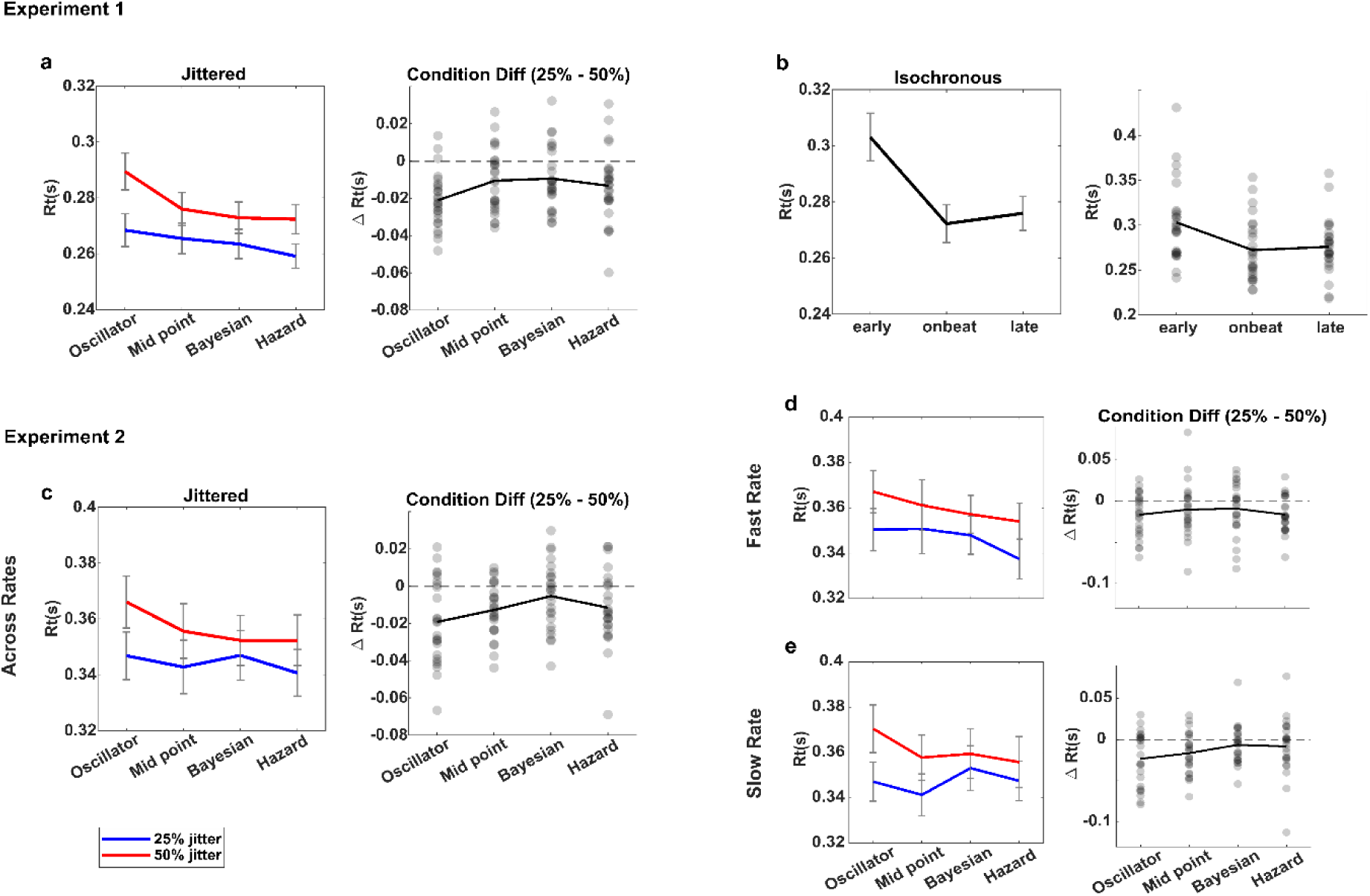
Behavioral performance in the speeded detection experiments, demonstrating temporal prediction without deterministic regularities. **a.** Experiment 1, single mean stream rate (1.25 Hz), jittered conditions. Left: Mean RTs for the target position conditions sorted according to the length of target interval in MV (25% jitter) and HV (50% jitter) conditions. Right: mean RT differences between 25% and 50% condition for each participant and target position; Each grey dot represents one participant, and the solid black line is connecting the mean RT differences across participants. **b.** Experiment 1, isochronous condition. Left: Mean RTs for the target position conditions. Right: mean RTs of each participant and target position; Each grey dot represents one participant, with solid black line connecting the group means. **c.** Experiment 2, two mean stream rates (1 Hz and 1.66 Hz). Same as a., across the two mean rate conditions. **d, e.** Same as c., separately for each mean rate (d: Fast, 1.66 Hz; e: Slow, 1 Hz) in both variability conditions.

For magnitude comparison and to verify ability to form Oscillator-based predictions we also tested an Isochronous condition (Fig. 5b). As typically observed, RTs were significantly faster at the on-beat time point compared to off-beat positions, t(25)=−5.28, p<.001, d=1.04. These results confirm that participants were able to intentionally drive oscillatory entrainment in deterministic environments. While the effect of on-vs. off-beat is not directly comparable to the uncertainty manipulation, note that as expected, the effect size in the isochronous condition was larger than the effect of Oscillator-based prediction in jittered streams.

### Oscillator-based predictions can be formed flexibly in varying environments independently of Bayesian learning

The goal of Experiment 2 (N=25) was to examine the flexibility of Oscillator-based temporal predictions in adapting to changes in the mean rate of the environment, while also excluding the possibility that the behavioral advantage observed in Experiment 1 reflected learning of the condition-specific ISI distributions rather than oscillator dynamics. Accordingly, we tested the same jittered conditions (MV: ISI SD=25%; HV: ISI SD=50%), but with two different mean rates of 1 Hz and 1.66 Hz (1000 and 600 ms mean interval, respectively) mixed within block, which were followed by the same target position conditions relative to the mean rate. If Oscillator-based prediction operates through flexible frequency tuning, behavioral effects should generalize across mean rates.

Aggregating across both mean rate conditions (Fig. 5c), like in Experiment 1, RTs were overall faster for MV compared to HV conditions (2×4 repeated-measures ANOVA, F(1,24)=30.54, p<.001, ηp²=.56), and varied between target positions (F(3,72)=4.55, p=.006, ηp²=.16). The interaction between jitter and target position was marginally significant, F(3,72)=2.63, p=.056, ηp²=.10, in line with differential effects of temporal variability across prediction points. Planned contrasts revealed that as in Experiment 1, RTs for Oscillator-predicted targets were faster in the MV than in the HV condition, t(24)=−4.30, p<.001(one-tailed), d=.86. In contrast, Bayesian-predicted targets did not differ between jitter levels, t(24)=−1.50, p=.15, d=.30. To examine again whether this Oscillator-based advantage in the MV condition reflected model-specific prediction beyond a global speeding of responses, we repeated the analysis after correcting model-predicted targets relative to the hazard control target. This revealed a weaker though marginally significant RT reduction (t(24)=−1.57, p=.065(one-tailed), d=.31. Finally, similar to Experiment 1, directly comparing the impact of reduced jitter on the Oscillator-and Bayesian-predicted targets revealed a stronger RT benefit for Oscillator-based targets (2×2 repeated-measures ANOVA, Jitter×Model interaction: (F(1,24)=5.96, p=.022, ηp²=.20). There was also a main effect of prediction model (F(1,24)=6.97, p=.014, ηp²=.22) but not of jitter (F(1,24)=.03, p=.86, ηp²=.001). Thus, despite the weaker effects, these results again show that Oscillator-based prediction operates in aperiodic streams when temporal variability is reduced, decoupled from Bayesian-based predictions, consistent with the pattern observed in Experiment 1.

Finally, we examined how Oscillator-based temporal predictions was affected by stimulus rates. A 2×2 repeated-measures ANOVA revealed faster RTs in the MV compared to HV condition (main effect of jitter: F(1,24)=17.26, p<.001, ηp²=.42), but neither a main effect of mean rate, F(1,24)<.001, p=.99, ηp²<.001, nor a Rate×Jitter interaction, F(1,24)=.91, p=.35, ηp²=.04. Together, these findings suggest that the behavioral advantage of Oscillator-based predictions is expressed similarly across different mean rates, supporting flexible rate tuning of the underlying oscillatory mechanism, while suggesting hazard-related response dynamics contribute substantially to the remaining between-condition differences.

## Discussion

Our environment often consists of continuous streams of events that are temporally jittered, devoid of deterministic temporal regularities that are typically considered a prerequisite for prediction. The present study reveals that even in such streams, an entrained oscillator mechanism, that is decoupled from Bayesian sequential updating, still operates to drive temporal prediction. Our model simulations show that while oscillator and Bayesian mechanisms behave similarly in certain conditions, increasing temporal variability made the model predictions more likely to diverge, with the Oscillator model being able to provide robust prediction even for streams with substantial jitter. The uncertainty representations of the models provided insight into the basis of this divergence, revealing greater sensitivity to local regularities in the Oscillator model vs. global statistics in the Bayesian model. Behavioral experiments using model-dissociating streams uncovered that, strikingly, humans demonstrated facilitated performance for Oscillator-predicted targets in 25% jittered streams, indicating that they were using this mechanism to form temporal predictions. In contrast, Bayesian predictions remained robust to temporal variability suggesting that they are less affected by the degree of temporal uncertainty. Together, these findings highlight that multiple computational strategies can coexist and contribute to temporal prediction in jittered environments, and establish oscillatory entrainment as a key component that enables tracking of transient local dynamics when these diverge from global regularities.

While aperiodic non-deterministic environments were traditionally considered ‘non-predictive’^2,14^, few recent studies suggested that oscillator or oscillator-approximating mechanisms with flexible frequency tuning guide prediction in constrained anisochronous environments^13,17^. Our study departs from previous investigations in significant ways. First, previous work focused on limited cases of low temporal jitter, or aperiodic streams constructed by jittering a latent isochronous stream. These encourage participants to infer an underlying beat or metrical structure^24,25^, but are also confounded with implicit arrhythmic regularities^26^. The present results, obtained without a latent rhythm and for up to 25% jitter, reveal the generalizability of oscillatory prediction in more ecological environments in which temporal regularities are local and transient rather than organized around a stable beat. Second, our results demonstrate that a fixed-period oscillator, a more parsimonious and biologically plausible mechanism^2,7,18^, is sufficient to explain temporal prediction in these contexts. This does not rule out the possible operation of an adaptive-frequency oscillator mechanism in aperiodic contexts, potentially as an implementation of Bayesian processes with which it overlaps^13,17^.

Our modelling analyses provided several key insights. First, they delineate the conditions for segregation between oscillator and Bayesian mechanisms. Oscillatory entrainment was shown to be mathematically equivalent to Bayesian estimation of phase and tempo^13,23^, or act as a hard-coded prior for a Bayesian inference process^17^. Consistent with this, our analyses show that the two mechanisms converge across multiple jittered environments, or under low temporal variability. However, they also show divergence of the predicted timepoints of these two mechanisms in specific jittered contexts as variability increases. Thus, Bayesian inference and rhythmic entrainment are not interchangeable descriptions of the same process, but can prioritize distinct aspects of temporal representation. Moreover, uncertainty trajectory and parameter-space sensitivity analyses showed that across streams with matched global statistics, Oscillator representation of uncertainty depended heavily on the specific arrangement of intervals, whereas Bayesian uncertainty was comparatively independent of specific realizations. Thus, the two mechanisms differ not only sporadically in their predicted timepoint, but fundamentally in the temporal scale at which information is represented, with Oscillator sensitive to the local temporal structure, whereas Bayesian to global distributional statistics^27,28^. Oscillator tracking of the environment would lead to a different predicted timepoint from Bayesian when local regularities transiently diverge from the global trend.

One interesting question is how phase concentration can be readout dynamically from an oscillatory mechanism. Instantaneous representation of uncertainty/confidence is essential for scaling prediction strength, supported by facilitated behavior with lower Bayesian uncertainty^29,30^ or higher oscillatory phase concentration^2,31^. In our simulations, we calculated phase concentration across different initial angles, conceptualized as derived across multiple trials. However, we propose that an equivalent computation could occur in a single-trial through a population of desynchronized period-matched oscillators, with confidence reflected in the degree of phase synchronization they demonstrate in response to input. This idea is consistent with evidence for coordinated oscillatory activity of different neural populations and neural oscillation modulations correlating with single-trial human behavior^7,16,32–35^. Moreover, a population view also provides a potential mechanism for encoding global temporal regularity. Our simulations show that even in jittered streams, oscillators whose natural frequency matched the mean input rate showed superior phase concentration and sustained activity, by this intrinsically representing the global mean rate (for similar multi-frequency oscillator array in isochronous streams see^16,22^). Under strong entrainment, however, this selectivity was reduced as even mismatched oscillators became partially synchronized, pointing to a potential trade-off between representing temporal rate and uncertainty, depending on entrainment strength.

Guided by the modelling results, the behavioral experiments examined whether oscillator and Bayesian mechanisms are in fact used for temporal prediction in aperiodic streams. The main finding was performance benefits at Oscillator-predicted targets in streams with 25% jitter, which enable stronger temporal prediction for both models due to higher prediction certainty, compared to 50% jitter. Critically, we used streams in which Oscillator-predicted target times were separated from Bayesian-predicted, and as such of adaptive-frequency oscillator^17^. This separation exceeded typical interval-discrimination thresholds^36^,making it unlikely that these intervals were integrated or confused with one another. Thus, these results reveal the psychological reality of a fixed-period entrained oscillator in guiding temporal prediction even under substantial input variability. Integrated with the modelling analyses, these findings suggest that in parallel to tracking global statistics, the brain continues to track local dynamics, which could be beneficial when these transiently diverge from the general trend. These patterns were replicated when the stream rate varied trial-by-trial (Experiment 2), pointing to the rapid switch between input-matching oscillators. This adds to similar findings of oscillatory entrainment to isochronous streams^37–39^, indicating that temporal prediction involves flexible engagement of oscillatory processes whose frequencies match the temporal structure currently present in the environment.

Bayesian mechanisms are well established in temporal learning and decision-making^10,40,41^. As such, the absence of selective benefit for Bayesian-predicted targets in the 25% compared to the 50% conditions should be interpreted cautiously. One possibility is that Bayesian mechanisms still operate, but integrate information not only within but also across trials. As in Experiment 1 the mean interval was identical across trials, this may have led to overall low uncertainty in both conditions, masking the influence of lower within-trial uncertainty in the 25% condition. Consistent with cross-trial integration, responses in Experiment 2 were numerically faster at target time points corresponding to the mean interval across rates (Fig. 5d,e for Hazard and Midpoint targets, respectively). Our simulations used a Bayesian model without a forgetting term, for parsimony and to capture within-trial accumulation of temporal information. However, this may not fully capture the potential updating across trials with different rates. Future work could extend this framework by testing alternative Bayesian architectures with integration of limited temporal memory^42^. Another possibility is that RT modulations for Bayesian-predicted targets were masked by the robust foreperiod effect we observed, i.e. increasingly faster responses to later targets^8,43^. Because Bayesian-predicted targets occurred later than Oscillator-predicted targets, they would have been more susceptible to such effect. At the same time, faster responses at matched control timepoints in the 25% condition indicate that participants were sensitive to the uncertainty structure of the stream, which is essentially a Bayesian process. Thus, these results should not be taken as evidence against Bayesian inference in aperiodic streams, and future work should evaluate this using more sensitive covert measures of neural encoding of Bayesian surprise^10,44^. Importantly, whether Bayesian mechanisms operate in a distinct timepoint or not is independent of the observed benefits at Oscillator-predicted targets.

The present findings also have broader implications for models of rhythm-based prediction. Oscillatory entrainment has been the mainstream explanation for predictions in rhythmic streams^2,7,45^, but this has been questioned due to suggestions that interval prediction mechanisms could operate sequentially, essentially leading to overlapping behavioral and neural modulations (see Fig. 2b for full overlap in isochronous streams)^16, 46–48^.The present study examines a case in which oscillator-based predictions are dissociated from interval-based predictions, derived from Bayesian inference rather than deterministically. The observed effects on Oscillator-predicted targets constitute the most direct evidence that rhythmic prediction cannot be fully reduced to interval-based prediction. This complements evidence of dissociations between rhythm-based and interval-based timing mechanisms^47,49,50^, suggesting that computationally distinct mechanisms may nevertheless converge on overlapping behavioral and neural patterns. Future work should combine computational modeling with neural recordings, to uncover how these processes are supported by parallel, converging, or dynamically weighted neural processes in naturalistic continuous environments.

Together, our findings point to the parallel operation of multiple predictive processes, based on local temporal dynamics, global statistical parameters, and the passage of time, whose relative contributions are shaped by environmental variability and task structure. Importantly, they suggest that temporal prediction do not need to rely on a single computational principle, but may emerge from the flexible interplay of distinct processes. In ecological settings, where temporal structure is neither strictly rhythmic nor entirely random but varies across timescales, such flexibility may be critical for adaptive behavior. By dissociating predictive principles under highly variable conditions, our study provides a first step towards understanding non-deterministic temporal prediction in fully naturalistic environments.

## Methods

### Computational Modelling

We implemented two computational models to investigate how temporal predictions could form in environments without deterministic temporal regularities. The first was a fixed-period coupled oscillator model, which describes how endogenous neural oscillations can entrain to external streams. The second was a Bayesian sequential updating model, which describes how current evidence is integrated with prior beliefs about the temporal structure of the environment. Both models provide time-resolved estimates of temporal prediction and associated uncertainty. In the Oscillator model these dynamics arise from the continuous evolution of oscillator phase, whereas in the Bayesian model they are updated sequentially following each observed interval.

### Coupled Oscillator Model

The oscillatory entrainment mechanism was modelled using a canonical nonlinear oscillating dynamical system, a biophysically plausible model widely used to simulate neural entrainment to rhythmic stimuli^22^. This framework assumes that an oscillator reflects fluctuations between high and low excitability states, whose phase can be coupled to the events of the environment. This alignment drives temporal prediction through the oscillator’s self-sustaining limit-cycle dynamics, leading to the next optimal phase occurring at a specific time point in the future. This modelling approach is commonly employed in the neural resonance framework of rhythm perception^22,51^. The dynamical system was characterized by two state variables, the radius (*r*) and angle (*θ*). This system produces the time-resolved dynamics of a canonical oscillator given some initial state (*r*, *θ*) and the time-resolved state of the environment s(*t*) (time-varying stimulation):

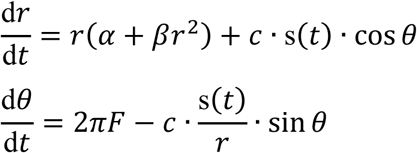

In this formulation, the α (bifurcation) and β (nonlinear saturation) parameters control the amplitude dynamics. These parameters were set to 0.5 and-2, respectively, leading to the system producing a self-sustaining oscillation at frequency *F* (in Hz) at a radius of *r* = 0.5. The additional parameter of this model, *c*, controls the strength of coupling to external inputs (scaled such that 1 would lead to a strong correction from 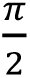 to 0 in one ms independently of sampling rate). When s(*t*) ≠ 0, the external input leads to a shift in the oscillator’s phase towards *θ* = 0, which is here assumed to be the optimal phase. In isochronous streams, this process leads to gradual alignment of the oscillator’s optimal phase with on-beat time, referred to as entrainment^2,16,22^. The value of c was set such that the model would lead to a similar magnitude of phase concentration (PC) increase within four isochronous stimuli across initial angles as was observed in previous work with four events^52^. Importantly, this model formulation did not contain a frequency detuning term^22^ making it a fixed-frequency oscillator, that is, external inputs can only lead to phase and amplitude adjustments. Based on this model, temporal prediction is defined relative to the last input as the next time in which the optimal phase is reached.

### Bayesian Sequential Updating Model

Temporal predictions based on learning the distribution of the inter-stimulus intervals (ISIs) were modelled using a sequential Bayesian updating framework^53,54^. In this framework, each newly observed sensory information, here the most recent ISI, is integrated with existing (Prior) knowledge to update the beliefs about the environment (Posterior), and hence about the next ISI. Rather than using a simple ideal Bayesian observer model^10^, we used a conjugate normal–gamma prior formulation^55,56^, which enables estimating both the mean and precision of a distribution, respectively. The model assumes that interval durations are drawn from a normal distribution with unknown mean and precision, for which a normal–gamma distribution serves as the conjugate prior:

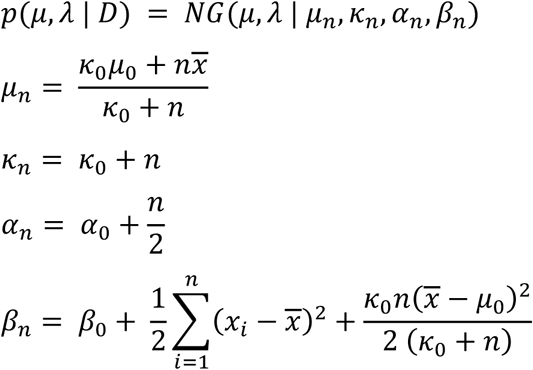

The two key parameters being estimated in this model are μ, the mean of the normal distribution (representing the predicted interval duration) and λ, the precision parameter reflecting prediction certainty. The other parameters of the model are κ, the effective number of observations contributing to the estimate; and α and β, the shape and scale parameters of the gamma distribution over precision. The ratio between the scale and shape parameters determines the expected uncertainty around the interval estimate (λₙ = βₙ(κₙ + 1)/(αₙκₙ)). Because the model updates its beliefs sequentially after each observed interval, the number of new observations at each step was n = 1, and the estimates of all parameters were updated after each ISI. In all simulations, we assumed a non-informative prior over interval duration at the beginning of each stream. Accordingly, the initial mean μ₀ was set to be uninformative and the prior observation weight κ₀ was set close to zero, minimizing the influence of the initial prior relative to the incoming observations. Based on this model, temporal prediction for the next interval after each ISI is defined as the posterior mean of the estimated normal distribution, whereas the posterior mean of the estimated inverse-gamma distribution characterized the associated prediction uncertainty. Notably, in this formulation, the model retains full memory of all previously observed intervals during sequential updating. Under this assumption, the final prediction for each stream corresponds to the mean of all observed intervals of that specific stream.

### Model Simulations

We conducted a series of simulations of the two models to study their behavior in environments with varying degrees of ISI variability, across a range of model parameters. The environment was modelled as a stimulus stream consisting of a sequence of brief sensory inputs. For simplicity we assumed that the stimuli can be characterized by their onset times, and the interval between the onsets of two successive stimuli was considered as the ISI. Stimulus streams consisted of sequences of 5, 7, 9 or 11 ISIs. All the streams had a fixed mean ISI of 800 ms but could have different levels of variability (SD range = 15%, 25%, 30%, 40%, 50%, 70%). To create a stream with length n and a given SD, we generated a large pool of candidate ISIs from a gaussian distribution with this SD and a mean of 800, and then systematically subsampled from this pool by sorting values in ascending order and selecting values at regular intervals of every n-th percentile across this distribution. This systematic subsampling ensured a balanced coverage of the distribution within each stream, and that the resulting stream would have the desired mean and SD. This approach was preferred over other previously used methods to generate jittered streams, such as independent random sampling of each interval, which can lead to individual streams having different mean, SD and coverage due to sampling variability, and jittering the ISIs around the intervals of an isochronous stream, which was shown to create correlation structures that can be tracked by humans^57^. To obtain a pool of different streams in each stream length and ISI SD combination, we randomly shuffled the set of ISIs within the constructed stream of that combination. All simulated streams were used as inputs to both models, and their prediction and uncertainty dynamics were tracked over time.

To simulate model behavior for a given stream, the Bayesian model was inputted the ISIs sequentially as they would have been experience in the continuous environment, with the parameters updating after each ISI. To account for any potential biases due to the initial prior selection, we ran the Bayesian model with different initial prior values (equally spaced around the range of ISI distribution of the stream) and an arbitrarily chosen shape and scale parameters for the initial uncertainty around the prior values, and the results were averaged across repetitions. For the Oscillator model, the ISIs of a given stream were converted into a time-series with a sampling rate of 1000 Hz of stimulus energy (1=stimulus onset, 0 otherwise), with the intervals between stimulus onsets matching the ISIs sequence. To simulate the dynamics of this system for a given [*θ*, *r*] initial state, we numerically integrated it using a fourth order Runge-Kutta algorithm with a fixed rather than variable integration step (MATLAB function ode4.m), to verify not skipping events in the stream. To obtain the prediction and uncertainty of the Oscillator model we ran the model 80 times, each time with a randomly chosen initial phase angle between 0 and 2pi. We computed the circular mean phase across iterations to track model’s temporal prediction (with *θ_optimal_* = 0, the optimal phase, being the peak of prediction), and quantified prediction certainty as the increase in PC relative to the initial state before the stream, that is, the PC of the randomly selected initial angles. PC was calculated as the mean vector length using the 80 angles at each time point using:

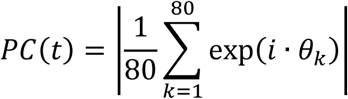

To overcome potential biases due to the random selection of the initial angles, for each stimulus stream this process was repeated 100 times, each with a new random set of initial angles, and the results were averaged across repetitions. As a measure of uncertainty we used the circular variance, defined as Phase Variance (PV); calculated as *PV = 1-PC* ^58^.

To examine the potential overlap or divergence between these two models in isochronous and jittered streams, we compared the two models in multiple levels. First, we compared their final predicted interval after exposure to all ISIs of a given stream. Second, we compared their uncertainty level in this prediction of the final interval. As the measure of prediction uncertainty is different between the two models (PV vs. posterior mean of inverse-gamma distribution), we transformed both measure into the change of uncertainty scaled to the uncertainty of a uniform distribution (circular or linear, respectively), across a range of initial states (angels for Oscillator, priors for Bayesian):

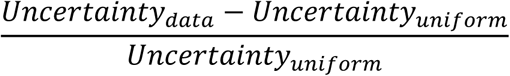

To assess whether the relationship between Oscillator and Bayesian uncertainty was linear or nonlinear, both linear and exponential regression models were fitted to the data for the isochronous and jittered conditions separately. Model comparison was performed using the Akaike Information Criterion (AIC), and the model with the lower AIC was considered to provide the better description of the data.

Third, we compared the dynamics of model uncertainties as the stimulus stream unfolds, to examine whether the final uncertainty around the predicted intervals might be achieved through different representational trajectories. Notably, we did not compare the dynamics of the predicted time points themselves as in the a priori chosen parameter regime the Oscillator model does not reliably converge to a stable PC during the early intervals of a jittered stream. Therefore, while circular averaging would still provide a mean that can be thought of as a “predicted point”, this averaging of noise is meaningless. Furthermore, as the Bayesian model is a conjugation of two distributions tracking mean and volatility of the environment, in this specific analysis, we calculated also the variance of the generated predictions from different initial priors (“Prediction Variance”) in addition to the model driven uncertainty governing inferred volatility. With this we opt to have different analogies to the oscillator PV which is considered as a measure of uncertainty around mean phase but can also be evaluated as the variance of different predicted timepoints (optimal phases) across multiple oscillators.

Fourth, we conducted an exhaustive investigation of parameter space, focusing on input sensitivity and forgetting, via analogous model-specific parameters. For the Oscillator model, we varied these through the entrainment strength (c) and dampening (through adjusting α & β), respectively. For the Bayesian model, to allow exploration of potential memory constraints, we introduced for this analysis only a memory parameter *mf*, defined on the interval [0,1]; κₙ = *mf*(*κ₀ + n*), that scales the effective confidence associated with accumulated observations while leaving the remaining Bayesian update equations unchanged. We then manipulated input sensitivity and forgetting through prior dependence (κ₀) and memory factor (*mf*), respectively. As outcome measures, we quantified the change in relative uncertainty of each model across their parameter values. Simulations were conducted for both isochronous and jittered (25% SD) streams to compare model behavior under deterministic and non-deterministic conditions.

Finally, to assess the specificity of oscillator recruitment, we tested whether oscillators with different intrinsic frequencies could entrain to input streams with a given mean rate. We compared PC dynamics across oscillators with varying natural frequencies (e.g., 0.8 Hz, 1.25 Hz, 2Hz) when exposed to isochronous 1.25 Hz streams under different entrainment strengths. We extended this analysis to jittered streams with a 1.25 Hz mean rate (15% and 25% SD), including both frequency-matched and mismatched oscillators (e.g., 0.8 Hz, 1.25 Hz, 2 Hz), to evaluate how frequency alignment and entrainment strength influence the selectivity and accuracy of temporal predictions.

## Behavioral Experiments

### Participants

A total of 51 participants took part in the study (Experiment 1: n = 26, 18 female, age range 22-31, mean age = 25.8; Experiment 2: n = 25, 15 female, age range 21-33, mean age = 25.6). All participants had normal or corrected-to-normal vision, and no history of neurological or psychiatric disorders. Informed consent was obtained from all participants in accordance with the ethical guidelines of Max-Planck Institute for Biological Cybernetics, and the study was approved by the Ethics Committee at the Medical Faculty of the Eberhard Karls University of Tübingen.

### Experimental Design and Stimuli

Tasks were implemented using the Psychtoolbox v.3.0 package for MATLAB^59^. In each trial, participants were presented with sequences of white visual stimuli occurring with variable ISIs, followed in most cases by a red visual stimulus (target) at the end of the stream. All the visual stimuli were circular patches with a diameter of 100 pixels, subtending approximately 1.8° of visual angle, with roughly 1-meter viewing distance (not head-fixed) and a resolution of 1024×768 pixels on a 17-inch CRT monitor (60 Hz refresh rate), presented foveally on a dark gray background.

The key experimental factor was the degree of ISI variability in the stimulus stream, which was either Medium Variability (MV, SD=25% of the mean) or High Variability (HV, SD=50%), manipulated in separate blocks. Critically, the model simulation results were used to guide the selection of the ISIs in each stream. Specifically, for each stream length and degree of variability, we selected from the set of simulated streams for this combination, a stream that maximized the divergence between model predictions, while also satisfying a minimum PC criterion in the Oscillator model under 25% ISI variability (phase-locking value > 0.2). In general, these selected streams led to the predicted target interval as the mean of the stream for the Bayesian model, and 0.5*this mean for the Oscillator model (the furthest point, circularly, from the prediction of the Bayesian model), with stronger certainty in both models for the 25% relative to the 50% condition. Importantly, we verified that to achieve this, the streams were not characterized by second-order regularities such as speeding up or slowing down.

In the first experiment streams could have a length of either 5,7,9 or 11 ISIs (equal probability), all with a mean of 800 ms (SD = 200 ms and 400 ms, in the LV and HV conditions, respectively), In the MV and HV blocks, each block consisted of 44 trials, 73% of them (32 out of 44) containing the red target stimulus. The key within-block manipulated factor was the time interval between the last stimulus of the stream and the target, which was set such that the target would appear at one of four different time points relative to the end of the stream. These time points could be based on Oscillator model prediction (∼ 406 ms, std = 44 ms), depending on precise timing in individual streams), Bayesian model prediction (800 ms), a time point in between the two model predictions (Midpoint, the mean of the Oscillator and Bayesian model predictions), and a late target to control for the hazard effect, i.e. increase in preparation with elapsed time (calculated based on Bayesian prediction uncertainty (i.e. 4*SD), ∼ 1064 ms, std = 48 ms). Each target type had the same probability of occurrence (8 out of 44 in each block), orthogonally to the length of the stream, such that every target interval appeared twice for each length. The remaining 12 out of 44 trials in each block (∼27%) were catch trials in which no target was presented (3 trials per stream length). The order of trials within a block was randomized.

In the first experiment only, participants also performed in separate blocks an isochronous condition, to verify that participants could indeed form a rhythm-based prediction and compare the magnitude of validity effect. In this condition, isochronous stimulus streams were constructed with a fixed ISI of 800 ms to match the mean rate of the jittered blocks, and could have a length of 5, 7, 9, or 11 ISIs with equal probability, matched to the jittered conditions. Each block of this condition consisted of 36 trials. For each stream length, targets could appear at three time points relative to the last stimulus of the stream: an early target (400 ms, 1 trial), an on-beat target aligned with the expected interval (800 ms, 4 trials), and a late target (1200 ms, 1 trial), along with catch trials (1000 ms of no stimulus, 3 trials) in which no target was presented. Thus, within each block, in 44% of the trials’ targets were on the beat and in 22% they were off the beat (the rest being catch trials, a ratio that was shown to be sufficient to drive synchronization to the beat and on-beat prediction. Trial order within each block was randomized. Isochronous blocks were presented at the end of the experiment to minimize potential carryover effects of entrainment from regular to variable streams. Sequence of trials in all blocks were constrained to limit immediate repetitions of exact same combination of ISIs and target intervals. Participants were instructed to respond as quickly as possible upon detecting the target stimulus and RTs to the targets were recorded.

The second experiment followed the general design of the first experiment, with few modifications. First, the experiment consisted of only MV and HV blocks (total of 8 blocks), without isochronous blocks. Second, each block consisted of 33 trials, 24 of which had a length of 11 ISIs (with 2 different ISI sequences, each a different shuffle of the ISI distribution), a change conducted to focus on long trials which could provide more PC in the Oscillator model. The other 9 trials were ‘filler’ trials with different lengths (ISI numbers: 5,7,13, equal probability), to prevent making the target fully predictable after 11 ISIs, and were not further analyzed. Most importantly, orthogonal to the block type and target position, each trial in each block had (with equal probability) one of two mean rates, fast or slow. With the same procedure as described in *Model Simulations*, we created pools of ISIs for the new rate conditions; in the fast condition the ISIs of the streams were drawn from a gaussian distribution with a mean of 600 ms (SD = 150 ms or 300 ms in MV or HV conditions, respectively) and in the slow condition of 1000 ms (SD = 250 ms or 500 ms in MV or HV variability conditions, respectively). The filler trials were created by scaling the ISIs values of the Experiment 1(μ = 800) to generate two distinct subsets with mean values shifted to μ₁ = 600 and μ₂ = 1000, respectively. Similar to the Experiment 1, the ISIs in the streams were selected based on the model simulations to maximize the differentiation between the predicted interval of the Oscillator and Bayesian models. In 67% of the trials in each block (16 out of 24), a target stimulus appeared at one of four different time points relative to the end of the stream. Similar to the first experiment, these time points could be based on Oscillator model prediction, Bayesian model prediction, a time point in between the two model predictions (Midpoint), and a late target to control for hazard. For the fast rate these times were ∼300 ms (std = 20 ms), ∼450 ms, 600 ms and ∼862 ms (std=49ms), respectively, whereas for the slow rate these were ∼482 ms (std = 55 ms), ∼742 ms, 1000 ms and ∼1300 ms (std = 71 ms), respectively. Each target type had the same probability of occurrence. In each block 8 out of 24 trials (∼33%) were catch trials in which no target was presented (1000 ms of empty window). In the 9 filler trials, the target ISI was randomly chosen out of the 4 possible positions while keeping their target probability the same across all blocks. The trials of each block were presented in randomized order. Participants were instructed to respond as quickly as possible upon detecting the target stimulus. Reaction times to the target were recorded.

### Procedure

The experiment was conducted in a dimly lit, acoustically isolated room. Participants received general instructions about the task but did not perform any practice trials, to prevent forming any explicit priors. In experiment 1, participants conducted 4 blocks of each of the MV and HV conditions, in alternating order, followed by 3 blocks of the isochronous condition. In experiment 2, participants conducted 4 blocks of each of the MV and HV conditions, in alternating order.

### Data Analysis and Statistics

Outlier rejection was performed by calculating Z-scores of reaction times (RTs) within each combination of stream variability and target position condition. Trials exceeding ±2.5 SD were excluded. In Experiment 1, this was applied across all stream lengths, whereas in Experiment 2, outlier rejection was applied within each rate condition (1 Hz and 1.66 Hz) for streams of 11 ISIs. The RTs were then averaged for each participant separately for each combination of stream variability and target position, and analyzed using repeated-measures ANOVAs and paired comparisons. For both experiments, a two-way repeated-measures ANOVA with Variability (medium, high) and Target Type (Oscillator-predicted, midpoint, Bayesian-predicted, hazard) as within-subject factors was first conducted to assess the effects of temporal variability and target predictability. To test the hypotheses that the reduction in uncertainty in the MV condition would lead to decreased RTs relative to the HV condition, we conducted planned contrasts using paired t-tests. To directly compare the models, we used a planned two-way ANOVA restricted to Oscillator-and Bayesian-predicted targets.

In Experiment 2, which included streams with two different mean rates, we repeated the same analysis approach, collapsing across rates. In addition, following the results of Experiment 1, a planned analysis examined the influence of mean rate on RTs to targets appearing at the Oscillator-predicted position, by performing a 2 × 2 repeated-measures ANOVA with factors Rate (fast, slow) and Variability (medium, high), as well as paired t-tests within rate to ensure that any observed effects of temporal uncertainty were not confounded by differences in stimulus rate. All analyses were conducted in MATLAB, with RTs restricted to a valid response window of 0.1–1 s. Effect sizes are reported as Cohen’s d for t-tests and partial eta squared (η²) for ANOVAs.

## Acknowledgements

This research was supported by funding from the Max Planck Society (Max Planck Independent Research Group) to Assaf Breska.

